# Increased insula activity precedes the formation of subjective Gestalt

**DOI:** 10.1101/2022.01.13.476145

**Authors:** Marilena Wilding, Christof Körner, Anja Ischebeck, Natalia Zaretskaya

## Abstract

The constructive nature of human perception sometimes leads us to perceiving rather complex impressions from simple sensory input. Bistable stimuli give us a rare opportunity to study the neural mechanisms behind this process. Such stimuli can be visually interpreted as simple or as more complex on the basis of the same sensory input. Previous studies demonstrated increased activity in the superior parietal cortex when participants perceived an illusory Gestalt impression compared to a simpler interpretation of individual elements. Here we tested whether activity related to the illusory Gestalt can be detected not only during, but also prior to it, by examining the slow fluctuations of resting-state-fMRI activity before the stimulus onset. We presented 31 participants with a bistable motion stimulus, which can be perceived either as four moving dot pairs (local) or two moving illusory squares (global). This allowed us to isolate the specific neural mechanisms that accompany the experience of an illusion under matched sensory input. fMRI was used to measure brain activity in a sparse event-related design. We observed stronger IPS and putamen responses to the stimulus when participants perceived the global interpretation compared to local, confirming the previously reported role of these areas in perceptual grouping. Most importantly, we also observed that the global stimulus interpretation was preceded by an increased activity of the bilateral dorsal insula, which is known to process saliency and gate information for conscious access. Our data suggest an important role of the dorsal insula in shaping an internally generated illusory Gestalt percept.

## 1. Introduction

Efficient processing of external visual input requires specialized neural mechanisms for grouping individual elements into a holistic Gestalt. Without these basic operations we would identify vital parts of our environment as randomly adjacent elements, lacking coherent meaning and, as a consequence, be soon overwhelmed by the input load. Grouping deficits are at the core of a neurological disorder known as *simultanagnosia* (Bálint, 1909), but have also been reported in psychiatric disorders, such as autism spectrum disorder (ASD). Individuals with ASD are often unable to identify the overall meaning of related visual elements in a scene or an object (Carther-Krone et al., 2016). As a consequence, they are frequently confronted with high neural workload due to processing single features as individual entities, which can cause serious challenges in their everyday lives (Scherf et al., 2008). ASD therefore exemplified the importance of these basic grouping operations.

A special type of bistable Gestalt stimuli gives us a rare opportunity to study the neural mechanisms behind these grouping processes. In contrast to traditional bistable stimuli that have two equally complex perceptual interpretations, these stimuli can be visually interpreted either as a collection of local elements or as one whole Gestalt on the basis of the same sensory input (Anstis & Kim, 2011; Lorenceau & Shiffrar, 1992). Identical sensory information allows for a comparison between the brain states during the grouped and the ungrouped perception that is free from low-level sensory biases. Previous studies repeatedly showed increased activity in parts of the parietal cortex (especially IPS and supramarginal gyrus) and a decrease in early visual areas (V1, V2) when participants perceived ambiguous stimuli as a grouped illusory Gestalt rather than an array of single elements (Carther-Krone et al., 2020; De-Wit et al., 2012; Fang et al., 2008; Grassi et al., 2016; Murray et al., 2002; Zaretskaya et al., 2013). One study also found subcortical activity in the putamen during the grouped interpretation (Zaretskaya et al., 2013). But what exactly determines whether a Gestalt can be formed? Given the fact that Gestalt in bistable paradigms appears and disappears spontaneously in the absence of any sensory change (i.e., it is generated internally), its formation may be related to the spontaneous ongoing fluctuations of activity within the intrinsic brain networks (Damoiseaux et al., 2006; Fox et al., 2005; Heuvel & Pol, 2010; Palva & Palva, 2011; Rosazza & Minati, 2011).

Only a few fMRI investigations focused on the role of spontaneous activity in subjective perception, with two major findings. On the one hand, there is strong evidence for the role of spontaneous prestimulus activity in stimulus-selective brain regions. For example, in one of the studies using an ambiguous face-vase illusion, the authors found increased activity in the fusiform face area (FFA) before the actual stimulus onset when participants subsequently perceived the two faces instead of a vase (Hesselmann et al., 2008a). Similar effects were also shown in the motion area MT+ in a near-threshold coherence detection task (Hesselmann et al., 2008b). On the other hand, there is also evidence for the increased prestimulus activity in a salience and alertness network that includes the dorsal anterior insular cortex (dAIC) and anterior cingulate cortex (ACC) in near-threshold tasks (Sterzer & Kleinschmidt, 2010). Several studies found higher ongoing activity in the dAIC, ACC and the thalamus prior to successful auditory stimulus detection (Sadaghiani et al., 2009), prior to correct responses in a discrimination task (Sadaghiani & D’Esposito, 2015), and prior to increased response speed in an auditory and visual detection task (Coste & Kleinschmidt, 2016). These findings point to an important role of this network as indicator of the level of alertness, which can facilitate subsequent stimulus processing. Importantly, in line with the first group of findings, these studies also reported increased prestimulus activity in regions relevant for stimulus processing, such as the auditory and/or visual cortex (Coste & Kleinschmidt, 2016; Sadaghiani et al., 2009).

Here we investigated the prestimulus neural correlates of perceiving subjective Gestalt in Gestalt-processing areas as well as in the insula region. We did this to determine whether brain activity fluctuations occurring *before* an ambiguous grouping stimulus is shown, can be predictive of a successful subsequent Gestalt perception. We presented participants with a bistable motion stimulus, which could be interpreted either as four locally moving dot pairs (local) or as two rotating illusory squares (global) (Anstis & Kim, 2011), while controlling for eye movements and other confounds. We interleaved short stimulus displays with long inter-stimulus intervals. This procedure allowed us to isolate the neural activity that precedes illusory Gestalt perception.

## 2. Material and Methods

Preregistration of the study design and methods was done before the start of data collection on *aspredicted*.*org* (https://aspredicted.org/blind.php?x=2de9nx).

### 2.1 Participants

41 healthy volunteers were tested in this study. After excluding ten participants due to predefined perceptual criteria (see below), and one who did not complete the whole experiment, 31 healthy participants between 19 and 31 years (14 female, mean age = 23.61, *SD* = 3.40) were ultimately included in the analysis. Participants were all right-handed and had normal or corrected-to-normal visual acuity. Further they had no history of neurological, psychiatric, or cardiovascular diseases and were not taking any medication regularly. All subjects gave written informed consent and underwent a standardized instruction. The study was conducted in accordance with the Declaration of Helsinki and was approved by the local ethics committee of the University of Graz.

### 2.2 Stimulus presentation and procedure

The stimulus (see Figure 1A) was presented on a gamma-corrected MRI-compatible monitor (NNL, Nordic-NeuroLab, Bergen, Norway) located behind the scanner bore with a resolution of 1920 × 1080 mm and a refresh rate of 60 Hz. Participants viewed the stimulus through a mirror attached to the head coil at a distance of 140 cm. Four pairs of black dots presented at 100 % contrast were shown on a grey background (luminance 202.5 cd/m^2^) around a white central fixation dot. The single dots had a size of 0.65° v.a. and were moving circularly in-phase. The distance between the center of the dot pairs and the center of the screen was 4.0°. The distance between the single dots in a pair was set individually for each subject in order to achieve a similar proportion of local and global responses. The exact distance value for each subject was determined based on the results of a test run before the main experiment. According to Anstis and Kim (2011), larger distance leads to an increase in the proportion of global perception. Therefore, to determine the optimal distance for each subject, we estimated the point of subjective equality (PSE) using the QUEST toolbox (Watson & Pelli, 1983) and an experimental paradigm similar to the main experiment (see below), but with 40 trials and a jittered inter-trial interval of 2-5 s. The mean distance between the paired dots over participants was approximately equal to the diameter of one dot (M = 0.66°, *SD* = 0.24°, min = 0.30°, max = 1.21°). The paradigm was presented using Psychtoolbox (PTB 3.0.17; Kleiner et al., 2007) and Octave (version 4.2.2), running on a Dell XPS13 7390 laptop (Dell Technologies, Round Rock, USA) under Linux Ubuntu 18.04 LTS.

**Figure 1.**
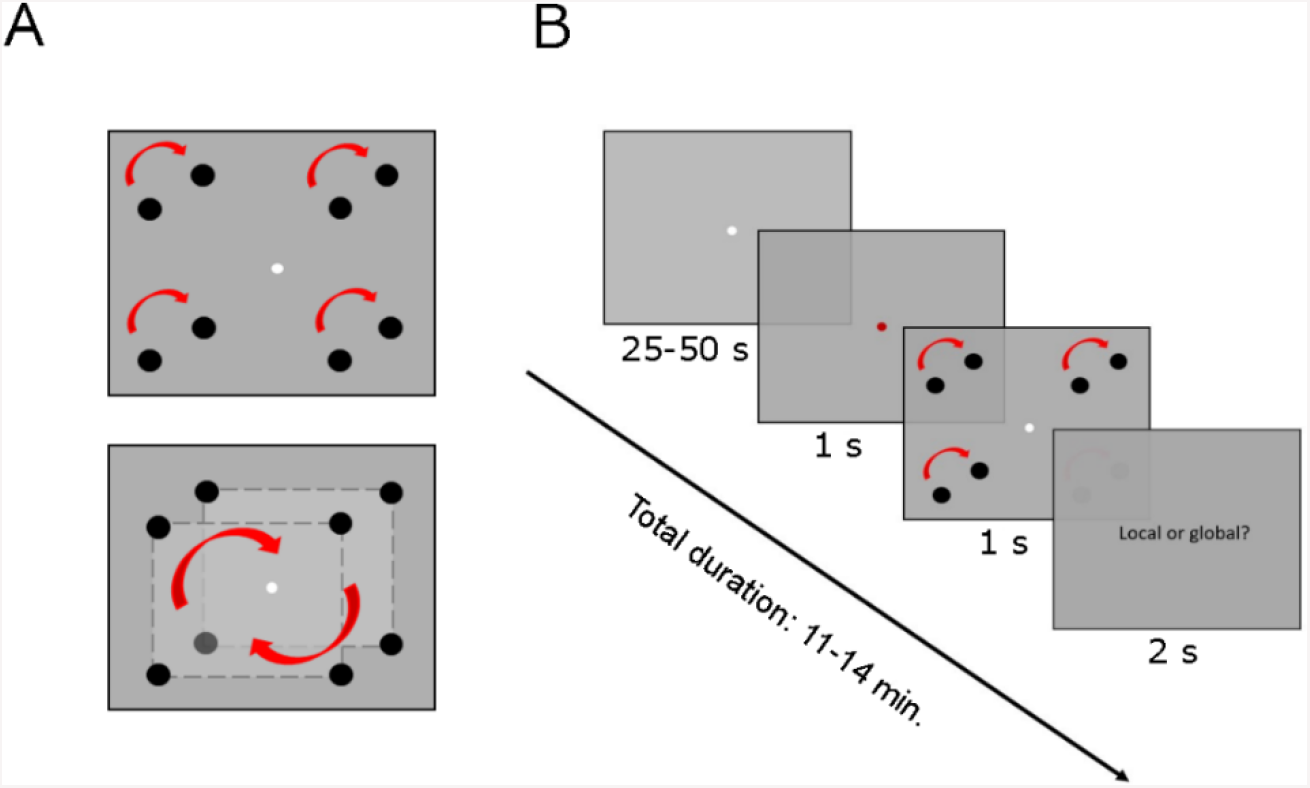
Illustration of the paradigm. A) Bistable motion stimulus with two possible stimulus interpretations: four locally moving dot pairs (top), or two rotating illusory squares (bottom) (Anstis & Kim, 2011). B) Depiction of one trial. After a jittered baseline period (25-50 s) showing a white fixation dot, the same dot turned red for one second, signalling stimulus onset. The stimulus was then also presented for one second. Subsequently, participants reported their spontaneous percept within two seconds after stimulus presentation via button press. They were required to fixate the dot in the center throughout the whole experiment. One run consisted of 18 trials and lasted for 11-14 minutes. Each participant underwent 6 runs, resulting in a total of 108 trials and approximately 75 minutes.

During the main experiment, participants were instructed to fixate the white dot in the center of the screen. After a jittered fixation period with durations ranging from 25 to 50 seconds with 5 s spacing (drawn from a gamma probability distribution), the white dot turned red for one second to signal the appearance of the upcoming stimulus, which then also lasted for one second. Participants were asked to indicate their spontaneously occurring percept by pressing two corresponding buttons on a button box in their right hand within two seconds after stimulus offset (see Figure 1B). The left button was used for local, the right for global percepts. Each run consisted of 18 trials and lasted between 11 and 14 minutes, depending on the exact duration of inter-stimulus intervals (mean of all runs and all subjects: *M* = 12.26 min, *SD* = 0.53). Each participant underwent six functional runs, resulting in a total measuring time of approximately 75 minutes (*M* = 73.57 min, *SD* = 1.16). The beginning of each run was time-locked to the image acquisition by a trigger sent from the scanner, which started the first trial at the start of the acquisition of the 5^th^ volume. In addition, to allow for clear separation of prestimulus activity from the stimulus onset, each baseline period ended with the nearest fully acquired MRI volume.

### 2.3 MRI Image Acquisition

MRI data were acquired on a 3 Tesla MRI system (Siemens Magnetom Vida, Erlangen, Germany) scanner using a 64-channel head coil. Functional images were obtained using blood oxygenation level dependent (BOLD) contrast with a simultaneous multislice echo planar imaging sequence (EPI) with the following parameters: repetition time (TR) = 880 ms, echo time (TE) = 30 ms, 45 interleaved acquired slices, multiband factor = 3, GRAPPA acceleration factor = 2, voxel size = 3×3 mm, no slice gap, flip angle = 65°, FOV = 210 mm. Additionally, a T1-weighted structural image was acquired for each participant with the following parameters: 192 slices, voxel size = 1 × 1 × 1 mm, TR = 1600 ms, TE = 2360 ms, TI = 1200 ms, flip angle = 9 °, FOV = 256 mm.

### 2.4 Physiological recordings

fMRI time course is typically contaminated by physiological noise sources like respiration and cardiac cycle. Modelling the physiological contribution to the fMRI signal in the analysis, especially in studies with prolonged periods without any task such as the resting state, has been shown to increase the contrast-to-noise ratio in fMRI experiments (Chang et al., 2009; Deckers et al., 2006; Verstynen & Deshpande, 2011). Therefore, physiological measures were recorded throughout the whole session using the standard hardware and software provided with the Siemens Vida scanner. Heart rate was recorded at a sampling rate of 400 Hz using a photoplethysmograph clip at the participants’ left index finger. Respiration was captured using a respiratory belt around their diaphragm, also with a sampling rate of 400 Hz.

### 2.5 Eyetracking

Due to prolonged fixation periods without any task that are essential for our experimental design, we wanted to make sure that our results can’t be explained by differences in fixation accuracy between conditions. Therefore, we performed eyetracking throughout the whole experiment. We used the MRI-compatible camera-based infrared Eyelink 1000 (SR Research, Ontario, Canada) eye tracking system to track position and pupil size of the left eye of each participant at a sampling rate of 500 Hz. Before the start of each run, participants underwent a 9-point calibration procedure to ensure a satisfactory tracking accuracy (spatial resolution of 0.50° or better). Eye tracking and data recording were controlled by a stimulus computer connected to the eye tracking PC via an Ethernet link. Two participants’ eyes were not trackable and were hence excluded from the following analysis of the eye tracking data (but not from the main fMRI analysis).

### 2.6 Post-experiment debriefing

After the main experiment, participants filled out a short questionnaire about their subjective experience during the task. This helped us to ensure a clean assignment of trials to the two conditions. Specifically, they were asked to indicate whether their subjective percept had changed within a single stimulus presentation period and whether they had corrected their perceptual choice within the response time window (both on a 5-point Likert scale, 1 = don’t agree at all, 5 = agree completely). If they had experienced more than one percept per trial, they were asked to report the number of such trials throughout the experiment. Participants who indicated a frequent change in responses (i.e., 5/5) and in perception (> 50% of trials) were excluded from the subsequent analysis (three participants in this study).

### 2.7 Data analysis

#### 2.7.1 Preprocessing of fMRI data

The functional and structural data were analyzed using the FreeSurfer and FS-FAST version 7.1.0 (http://surfer.nmr.mgh.harvard.edu; Dale et al., 1999; Fischl et al., 1999). The structural T1-weigted scan was processed using the FreeSurfer default recon-all pipeline and then visually inspected to insure correct segmentation and cortical surface reconstruction. Functional data were analyzed using the FreeSurfer’s functional analysis stream (FS-FAST). First, the initial four volumes of each functional run were excluded to allow for T1 equilibration effects. In the next step, the remaining volumes underwent motion correction, slice-timing correction and coregistration with the anatomical T1 scan using boundary-based registration (Greve & Fischl, 2009). The cortical data were then projected into the FreeSurfer fsaverage template surface space and smoothed using a 2D Gaussian kernel with the full-width at half-maximum (FWHM) of 6 mm. The subcortical data were mapped onto the MNI305 space and smoothed with a 3D Gaussian kernel with the FWHM of 6 mm.

#### 2.7.2 General linear model

Functional trials were assigned to one of the two conditions (either local or global) based on the behavioral responses. Trials without a response (on average 2%) were left out. The average BOLD signal time courses around each condition were extracted using the finite impulse response (FIR) GLM model with the time window -5.4 to 12.2 s relative to stimulus onset. With a repetition time of 0.88 s, the analysis thereby comprised 7 prestimulus timepoints and 14 poststimulus timepoints. The time window was chosen based on a previous fMRI study reporting prestimulus effects in the temporal lobe (Hesselmann et al., 2008a). In addition, scanner drifts and run-specific offsets were included as nuisance regressors. This FIR-based GLM allowed us to avoid any assumptions about the exact shape of pre-stimulus activity (albeit reducing the statistical power for detecting post-stimulus activity, which is typically assumed to have a canonical HRF shape).

To avoid the contamination of prestimulus resting-state activity by physiological signals, those were added to the GLM as further nuisance regressors. The physiological data were processed with the MATLAB-based *PhysIO toolbox* (Kasper et al., 2017). First, the data were aligned to the fMRI time series and preprocessed by recovering individual repetitive signal features from noise using a peak detection algorithm and discarding deficient data segments. Applying RETROICOR phase expansion (Glover et al., 2000), the periodic effects of pulsatile motion and field fluctuations were subsequently modelled as a Fourier expansion of cardiac and respiratory phase. The expansion orders were set to default, following the parameters of Harvey et al. (2008; 3^rd^ order cardiac model, 4^th^ order respiratory model, and 1^st^ order interaction model). The regressors were automatically downsampled to a reference slice of each volume and then added to the GLM as nuisance regressors. Three participants were lacking the physiological data due to technical issues while recording. For these participants, we used signals from the white matter and cerebrospinal fluid as nuisance regressors in the GLM instead.

#### 2.7.3 Regions of interest definition

To examine brain activity in regions known to be modulated by conscious illusory percepts in bistable paradigms (Grassi et al., 2018; Sadaghiani et al., 2009; Sterzer & Kleinschmidt, 2010; Zaretskaya et al., 2013) we conducted a regions of interest analysis. To avoid any potential biases due to analysis circularity (Kriegeskorte et al., 2009), regions of interests were defined using the following independent procedures. The anterior intraparietal sulcus (aIPS) region in each hemisphere was defined using a probabilistic atlas as the most anterior topographic map within the sulcus (IPS5; Wang et al., 2015). The insula and the putamen were defined using Freesurfer’s automatically generated parcellation (Desikan-Killiany and Destrieux atlas; Fischl et al., 2004) and segmentation atlas (Aseg atlas; Fischl et al., 2002). Early visual areas V1 and V2 were defined using V1 and V2 labels generated by FreeSurfer (Hinds et al., 2008).

#### 2.7.4 Region of interest analysis

To quantify the pre- and post-stimulus activity for each condition in each ROI, we averaged the individual-level beta estimates of the FIR model for each of the two conditions over all vertices (for the cortical regions) or voxels (for the subcortical regions) in a ROI. After this, ROI-level beta values belonging to the prestimulus time points were summed together to form one prestimulus net activity value per subject per condition. These values were used to compare prestimulus activity for the global and local conditions using a paired t-test. To confirm that we can replicate the results of the previously reported activity differences *after* the percept onset, the beta estimates of the central 6 poststimulus timepoints were summed together and processed similarly to the prestimulus data.

#### 2.7.5 Additional whole-brain analysis

To make sure we did not miss prominent prestimulus effects that were not part of our original hypothesis, we also conducted an explorative whole-brain analysis. This analysis followed the same procedure as the ROI analysis, but was conducted for every vertex on the surface and for every voxel of the subcortical structures. Beta values belonging to the prestimulus timepoints were summed together and the difference of sums for the global and local condition for every subject was used as a contrast estimate for the subsequent second-level group analysis. To test whether there is an overall prestimulus activity difference between conditions, we performed one-sample t-test with the individual-level contrast estimates. Results were corrected for multiple comparisons using the cluster correction method (Monte Carlo Simulation; Greve & Fischl, 2018) with a cluster-forming threshold of p <.05 (two-sided) and a cluster significance level of p <.05 (two-sided), with additional Bonferroni correction for the three spaces (two hemispheres and one subcortical space) of the cluster-level p-values.

#### 2.7.6 Eyetracker data analysis

The eyetracker data were processed in MATLAB R2019a (Mathworks, Natick, MA, USA). After linear interpolation of blink events in the gaze and pupil time courses, and additional z-scoring of the pupil data for each run (subtracting the mean and dividing by the standard deviation over time) the analysis was similar to the analysis of the functional MRI data. First, stimulus onset times were classified based on the participants’ behavioral report (local or global) and resampled to the fMRI time course by averaging all values within a time bin of ± TR/2 (0.44 s) around each volume acquisition onset. Prestimulus values for each condition (time points -5.4 to -0.12 s relative to stimulus onset) were summed and then compared in fixation accuracy (Euclidean distance between the eye position and the fixation dot) using a paired samples t-test. In addition, to rule out that spontaneous fluctuations in arousal can explain our fMRI results, we performed the same analysis using the pupil size as an indicator of arousal level (Nakano et al., 2021; Wang et al., 2018).

## 3. Results

### 3.1 Behavioral results

We first analyzed behavioral responses of our participants. The two interpretations of the stimulus were well balanced over subjects and runs (mean local: 53.3 %, *SD* = 8.9%; mean global: 46.6 %, *SD* = 8.9%; see Figure 2 illustrating the response distribution for each participant). The mean percentage of trials without a response (misses) over participants was 2.0 % (*SD* = 3.7%). There was no significant difference in the probability for two consecutive percepts to be different between local and global trials (average *p*_*local*_ = 60%, average *p*_*global*_ = 67 %; *t*_*30*_ = -1.0, *p* = 0.33, *d* = -0.18). Also, the time it took participants to indicate their percept did not differ significantly between conditions (mean_local_ = 0.83 s, ± SEM = 0.05; mean_global_ = 0.84 s, ± SEM = 0.05; *t*_*30*_ = - 0.51, *p* = 0.61, *d* = -0.09).

**Figure 2.**
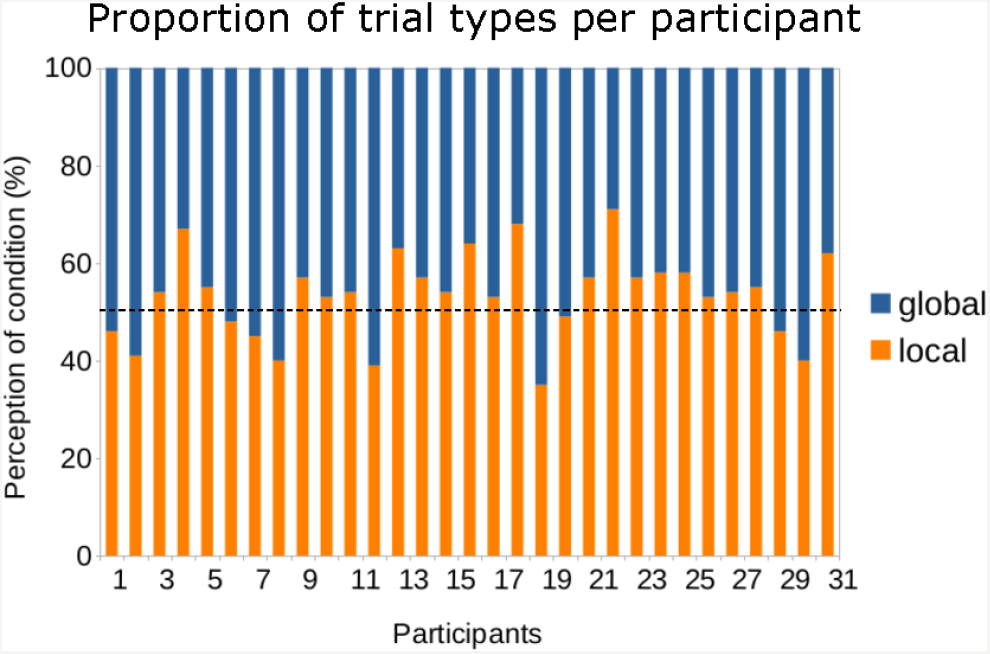
Proportion of global and local trials for each participant. In total, participants’ perception of both conditions was highly balanced (mean local: 53.3%, mean global: 46.6%). The black dotted line indicates 50% of trials, i.e., an optimal balance of percepts over runs.

### 3.2 fMRI results

#### 3.2.1 Regions of interest analysis

##### Poststimulus analysis

We first checked whether the percept-dependent effects found for this stimulus in a continuous bistable perception paradigm (i.e., activation of the IPS and putamen, deactivation of early visual areas during Gestalt perception) hold for our experimental design with short stimulus presentations and long inter-trial intervals. Indeed, we found that the left IPS responded stronger to the global compared to the local percept (*t*_*30*_ = 2.46, *p* = 0.02 *d* = 0.44). This difference was most pronounced at 5 to 7 s, consistent with the typical peak of the hemodynamic response function. Additionally, our data revealed stronger activity in the right putamen in response to a global percept (*t*_*30*_ = 2.36, *p* = 0.02, *d* = 0.42), see Figure 3A. The other ROIs didn’t show any significant difference in poststimulus activity between conditions (see Table 1 and Supplementary Figure S1).

**Table 1.** Values of the paired t-tests for each region of interest before (prestimulus) and after (poststimulus) stimulus onset.

| *Brain region* | *prestimulus* |  |  | *poststimulus* |  |  |
| --- | --- | --- | --- | --- | --- | --- |
| | $t_{30}$ | $p$ | $d$ | $t_{30}$ | $p$ | $d$ |
| Left IPS | 1.78 | 0.08 | 0.32 | **2.46** | **0.02** | **0.44** |
| Right IPS | 1.38 | 0.18 | 0.25 | 1.79 | 0.08 | 0.32 |
| Left insula | **2.23** | **0.03** | **0.40** | 1.30 | 0.20 | 0.23 |
| Right insula | **2.20** | **0.03** | **0.40** | 1.56 | 0.13 | 0.28 |
| Left V1 | 0.01 | 0.99 | 0.00 | -0.59 | 0.56 | -0.11 |
| Right V1 | -0.59 | 0.55 | -0.02 | -0.46 | 0.65 | -0.08 |
| Left V2 | -0.12 | 0.90 | -0.20 | -0.19 | 0.82 | -0.04 |
| Right V2 | -0.08 | 0.94 | -0.01 | -0.20 | 0.84 | -0.04 |
| Left putamen | 1.50 | 0.14 | 0.27 | 0.58 | 0.57 | 0.10 |
| Right putamen | 1.70 | 0.11 | 0.30 | **2.36** | **0.02** | **0.42** |

Bold values indicate a significant difference between conditions; $p < .05$ . Poststimulus values were derived from the 6 most central timepoints.

**Figure 3.**
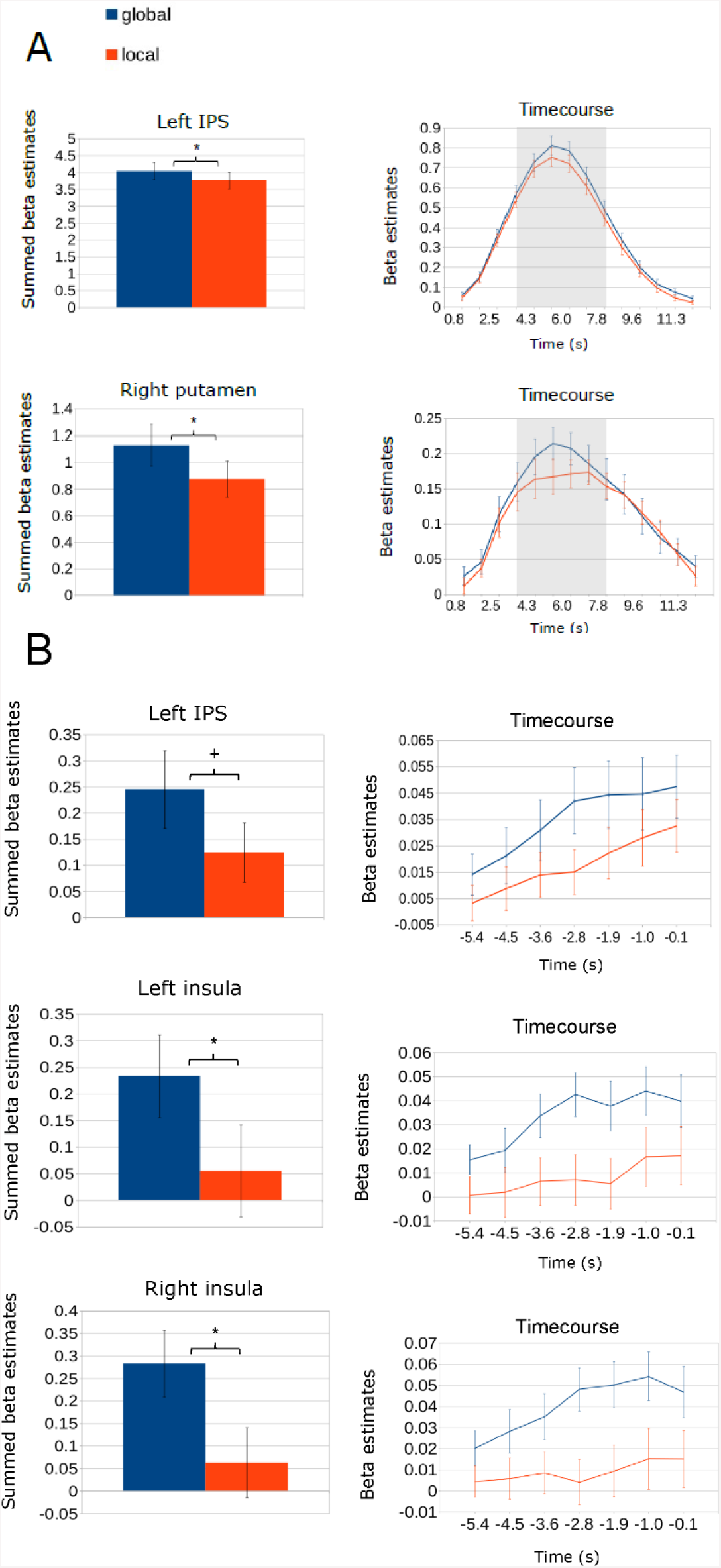
Average activity for both conditions for summed prestimulus (B) and poststimulus (A) timepoints in regions of interest. Prestimulus values were derived from the 7 timepoints of the FIR-GLM model within a time window of -5.4 to -0.12 s, relative to the stimulus onset. For poststimulus comparisons, the central 6 timepoints were summed (indicated by the grey area). A) In the left IPS, we found significantly stronger activity in global compared to local trials after stimulus onset, with a peak in activity at around 5-7 s. Additionally, we observed increased putamen activity after stimulus onset in global trials. B) Both the left and the right insula exhibited a significant difference in activation before stimulus onset. We also observed a trend towards significant difference between the conditions in the left IPS before stimulus onset. Error bars represent the SEM (±) over subjects. Significant differences are indicated by an asterisk (*p<.05). A trend (p<.08) is indicated by a “+” sign.

##### Prestimulus analysis

**Our primary goal was** to investigate whether specific regions of interest, which have been associated with perceptual grouping and conscious perception in prior studies, would also exhibit changes in activity *before* the actual stimulus presentation. Specifically, we hypothesized that the IPS, the putamen, the insula, and the early visual regions (V1, V2) would show not only poststimulus activity differences depending on the subjective perceptual content, but also a difference in *prestimulus* spontaneous brain activity fluctuations. We found that both the left and the right insula exhibited a significant difference in activation before stimulus onset (left insula: *t*_*30*_ = 2.23, *p* = 0.03, *d* = 0.40; right insula: *t*_*30*_ = 2.20, *p* = 0.03, *d* = 0.40), with the largest difference between conditions occurring between -2.8 s and -1.9 s for both hemispheres (see Figure 3B). In addition, a trend was observed in the left IPS (*t*_*30*_ = 1.78, *p* = 0.08, *d* = 0.32). The remaining regions of interest did not show any statistically significant difference in prestimulus activity between local and global trials (see Table 1 and Supplementary Figure S2).

Since the insula is comprised of several subparts with distinct functional roles (Kurth et al., 2010; Sterzer & Kleinschmidt, 2010; Uddin et al., 2017; Uddin, 2015), in a subsequent explorative analysis we further investigated which of those subparts (superior, central, inferior, and anterior) drive our result (see Figure 4). We found the bilateral superior and central part to be most engaged during global versus local trials before stimulus onset (superior left: *t*_*30*_ = 2.45, *p* = 0.02, *d* = 0.44; superior right: *t*_*30*_ = 2.87, *p* = 0.001, *d* = 0.52; central left: *t*_*30*_ = 2.25, *p* = 0.03, *d* = 0.40; central right: *t*_*30*_ = 2.31, *p* = 0.03, *d* = 0.42). In contrast, after stimulus onset, only the right central part was found to be more engaged in global trials (*t*_*30*_ = 2.52, *p* = 0.02, *d* = 0.45, all other poststimulus effects: *t*_*30*_ ≤0.85, *p* ≥0.40).

**Figure 4.**
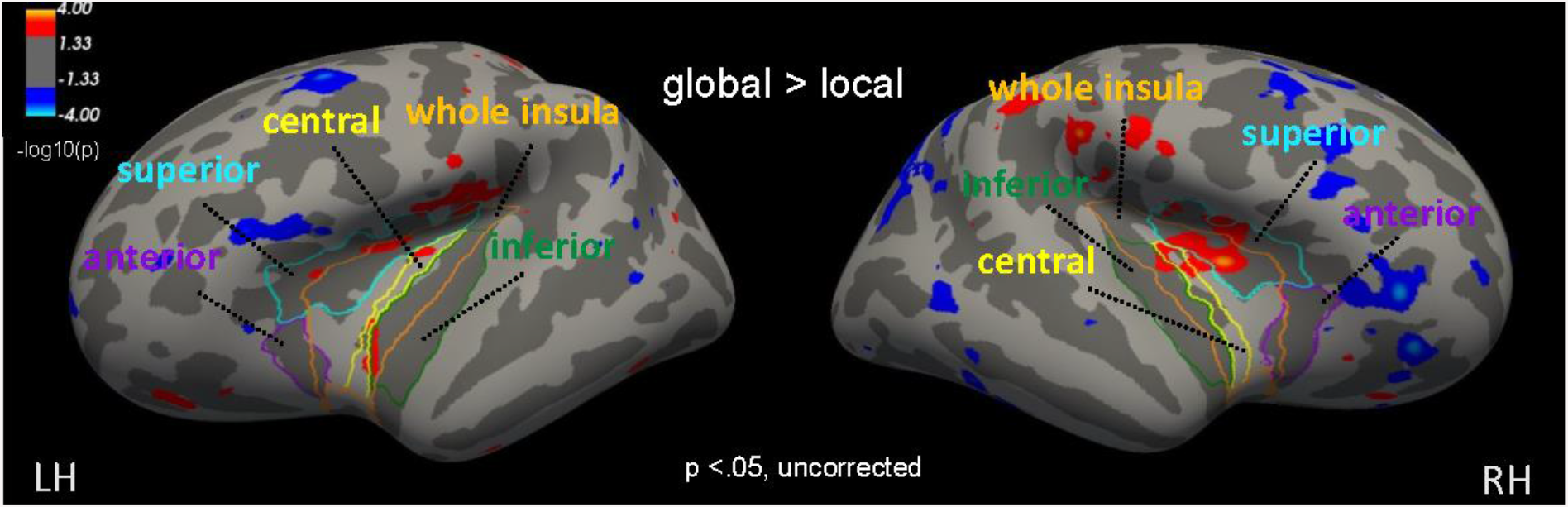
fMRI activity for the contrast global > local *before* stimulus onset (summed timepoints -5.4 to - 0.12 s) depicted on the inflated Freesurfer template cortical surface of the left and right hemisphere shown at p<.05 (uncorrected). Outlines show the cortical parcellation of the insula. LH, left hemisphere; RH, right hemisphere. Insula regions were defined using the Desikan-Killiany (whole insula) and the Destrieux (insula subparts) parcellation atlases provided by Freesurfer.

#### 3.2.2 Whole-brain analysis

To make sure our ROI analysis did not miss any brain areas that showed a significant prestimulus activity beyond the ones we had a specific hypothesis about, we additionally performed a whole-brain analysis. We did not find any significant clusters that survived the multiple comparisons correction, neither for the prestimulus nor for the poststimulus activity.

Given that our IPS ROI showed a trend towards significance before stimulus onset, in the second exploratory analysis we investigated the spatial correspondence between our parietal IPS ROI, the spatial distribution of the poststimulus activity, and the spatial distribution of the prestimulus activity within the intraparietal sulcus at p<.05 uncorrected. This analysis revealed only a partial overlap between the prestimulus activity and our IPS ROI shown in Figure 5, which could explain the statistical trend.

**Figure 5.**
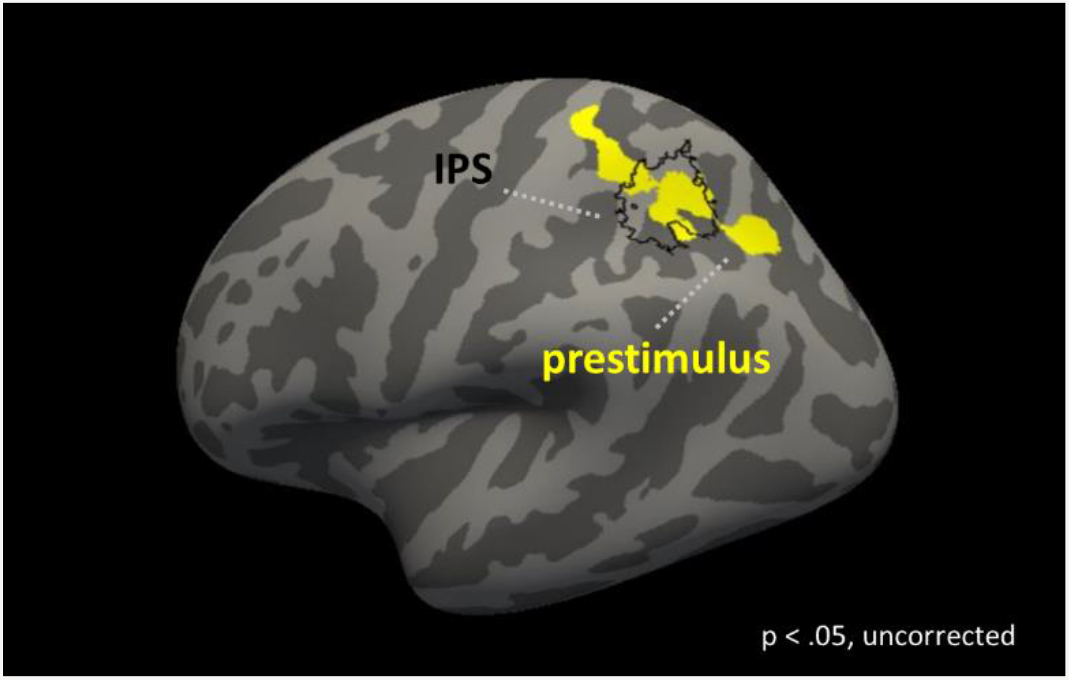
Spatial overlap between the IPS ROI location (black outline; corresponding to IPS5 from Wang et al., 2015) and prestimulus parietal activity (in yellow). Activity is depicted on the inflated Freesurfer fsaverage template cortical surface of the left hemisphere.

### 3.3 Eyetracking

To rule out any effect of prestimulus eye position on the participants’ perception, we computed a paired sample t-test for fixation accuracy, using the summed prestimulus timepoints. We did not find any significant differences in fixation accuracy between ‘local’ and ‘global’ trials before the stimulus onset (*t*_*28*_ = 1.09, *p* = 0.28, *d* = 0.20). Moreover, our data revealed no significant difference in prestimulus average pupil size between the two conditions (*t*_*28*_ = -0.21, *p* = 0.83, *d* = -0.04. Figures 6A and B show the group average peristimulus time course for fixation accuracy and pupil size, respectively.

**Figure 6.**
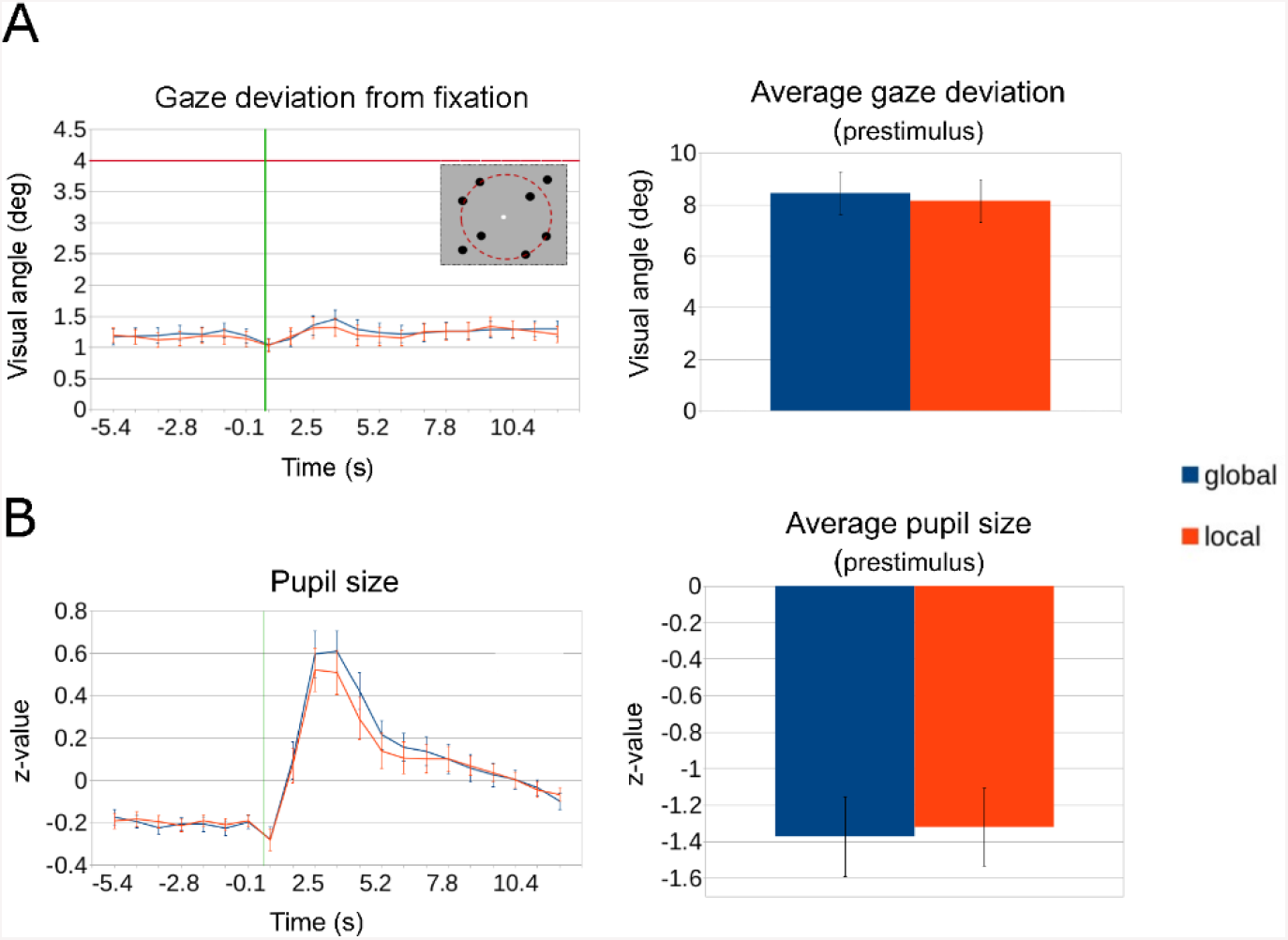
Eyetracking measures for global and local conditions. A) Average fixation accuracy (average Euclidean distance between the fixation dot and eye-position data). The red dotted horizontal line indicates the distance from the center of the screen to the center of a single dot pair in the stimulus (also indicated schematically on the inset at the top right). B) Average pupil size. The green line indicates stimulus onset. Error bars represent the SEM (±). Group analysis showed no significant difference between the local and the global condition before stimulus onset for both measures.

## 4. Discussion

The aim of this study was to identify possible neural correlates of Gestalt perception in spontaneous activity fluctuations *before* stimulus onset. Using a bistable stimulus with a global and a local interpretation of the same sensory input, we observed increased bilateral dorsal insula activity preceding trials that were interpreted as global. Overall, our results show that the bilateral insula seems to be responsible not only for gating sensory information for reaching conscious perception as reported previously, but also for the formation of a more complex illusory interpretation instead of a simpler interpretation of the identical consciously perceived sensory input.

### Salience/alertness and insula activity

Our results are in line with previous reports implicating increased prestimulus insula activity in subsequent improvement of different aspects of task performance. First, increased insula activity has been shown to predict successful stimulus detection. For example, Sadaghiani et al. (2009) used a near-threshold auditory detection task and found increased insular and auditory cortex activity prior to a successful auditory stimulus detection. Similar effects were also reported for face detection in the visual modality (Huang et al., 2021) as well as electrical stimulus detection in the somatosensory modality (Grund et al., 2021). Second, prestimulus insula activity was found to predict not only the success, but also the speed of stimulus detection. Coste & Kleinschmidt showed that faster detection in an auditory and a visual task is preceded by higher insular and visual cortex activity. Lastly, increased prestimulus insula activity was associated with increased performance accuracy in a discrimination task (Sadaghiani & D’Esposito, 2015). These studies speculated that the salience/alertness network involving the insula facilitates information processing by rendering an individual more receptive to sensory input (Sadaghiani et al., 2009; Sterzer & Kleinschmidt, 2010), or even by *gating* the information for conscious access (Huang et al., 2021).

There is a substantial difference between these studies and our current approach. While in the near-threshold detection paradigm sensory information either succeeds or fails to reach the conscious level, in our bistable stimulus paradigm the same consciously perceived information can either be formed into an illusory percept or not. In other words, illusory Gestalt formation occurs far beyond the level of conscious access. Our findings therefore suggest that a similar mechanism involving the salience/alertness network may operate in both cases. In a detection paradigm it may contribute to a more efficient processing of the sensory input, whereas in our bistable grouping paradigm to generating a more complex interpretation of the sensory input. Indeed, higher alertness has repeatedly been found to enhance visual grouping of single elements (Schneider, 2018; Weinbach & Henik, 2011), supporting this idea.

The human insula is a highly heterogenous brain region with several functional subdivisions (Uddin et al., 2017). Interestingly, the anterior insular cortex (AIC) activity has repeatedly been related to the presence of a stimulus, the recognition of its identity, as well as to stimulus changes (Dehaene et al., 2001; Ploran et al., 2007; Sterzer & Kleinschmidt, 2010). Since this region has been found relevant for a multitude of functions, including bistable paradigms (Knapen et al., 2011; Lumer et al., 1998), Chang et al. (2012) suggested a functional division into a dorsal anterior and a ventral anterior part. Importantly, the dorsal AIC is thought to be involved in higher-level cognitive operations, whereas the ventral AIC is associated with affective processes.

Our results are well in line with these findings. Since our main analysis did not differentiate between insula subregions, we conducted an additional explorative analysis for each of the insula subregions following the standard insula parcellation scheme (Fischl et al., 2004). This analysis showed the strongest prestimulus effect to be in the dorsal insula (see Figure 4), suggesting the specificity of our insula effect to its ‘cognitive’ part.

### Alertness versus arousal

When interpreting our insula results, it is crucial to note that alertness specifically refers to a cognitive state accompanied by increased attention and task-readiness. This should not be confused with arousal, which is associated with activity of the sympathetic nervous system and manifests itself primarily in physiological measures, such as pupil size and heart rate (Brown & Bowman, 2002). Although arousal is mainly expressed physiologically, it can still co-occur with a more attentive state. Here, a possible arousal-related influence on subjective stimulus interpretation was ruled out by two approaches. First, we analyzed pupil size data showing a similar pupil size for both conditions during the prestimulus time range. Second, we used physiological recordings of pulse and respiration to account for potential physiological confounds of fMRI data by including them into the general linear model. Thus, our results are unlikely to be due to arousal-related effects.

### Content-selective prestimulus activity

Previous investigations of prestimulus activity in bistable perception paradigms repeatedly emphasized the role of percept-selective areas in shaping the subsequent percept. For example, an fMRI study using the Rubin’s face-vase illusion reported increased activity in the percept-selective Fusiform Face Area (FFA) prior to the occurrence of a face percept (Hesselmann et al., 2008a). Another study investigating prestimulus oscillatory signatures of the subsequent percept in MEG, found increased synchronization between FFA and V1 when participants perceived a face, which was suggestive of feedback from the FFA to V1 (Rassi et al., 2019). Somewhat consistent with these findings, our results suggest a Gestalt-specific prestimulus activity in the IPS. This effect did not reach significance in our main ROI analysis, showing up only as a trend, but the whole-brain results clearly show prestimulus IPS activity in the left IPS, partially overlapping with the ROI. The IPS has been repeatedly shown to play a role in the formation of subjective Gestalt impressions for such bistable stimuli (Grassi et al., 2016; Zaretskaya et al., 2013), and therefore can be viewed as a percept-selective region. Our results therefore extend the previous findings on bistable perception by demonstrating that the insula, alongside the percept-selective areas, plays a role in shaping subjective perception. Whether these areas act independently or in cooperation is yet to be determined by future studies.

Our remaining regions of interest did not show a significant prestimulus activity difference. However, we did find increased putamen activity in response to the global interpretation, replicating previous results (Zaretskaya et al., 2013). What surprised us was that we did not observe a reduction in activity in the early visual areas (V1, V2) after global Gestalt perception in our summed analysis, contrary to what was shown by previous studies (De-Wit et al., 2012; Fang et al., 2008; Murray et al., 2002; Zaretskaya et al., 2013). By looking at the timepoints individually, however, we did observe bilateral deactivations in both V1 and V2 in response to the global stimulus interpretation. Interestingly, the difference between stimulus interpretations started to emerge after the peak activity, at around 6 s in both hemispheres (see Supplementary Figure S2). Since deactivation happened only in the second half of the response, summing 6 poststimulus timepoints around the peak together may have cancelled this effect. Overall, however, response time courses in the visual cortex imply that relevant information about the perceptual content may be contained not only in the response amplitude, but also in its shape, which is overlooked in more conventional modelling of the hemodynamic response using canonical HRF.

### Comparison of temporal characteristics

Our FIR analysis allowed us to look not only at prestimulus activity as a whole, but also at every timepoint individually and hence determine the temporal evolution of activity in each region of interest. Interestingly, we found that activity in the brain areas which show activity differences already before the subsequent percept (bilateral insula, left IPS) exhibit a simultaneous peak at around -2.8 to -1.9 s prior to stimulus onset. This is partially comparable with previous studies, which reported somewhat later times of strongest activity in stimulus-selective areas (Coste & Kleinschmidt, 2016; Hesselmann et al., 2008a.; Sadaghiani et al., 2009) and insular regions (Coste & Kleinschmidt, 2016; Huang et al., 2021), namely between 1.5 s before stimulus onset and 0.0 s. We do not know the reasons for this discrepancy. However, given the slow nature of resting-state fMRI signal fluctuations and the autocorrelations inherent in the fMRI signal, effects found further away from the stimulus onset makes them less likely to be confounded by the stimulus activity, thereby only strengthening our findings.

When interpreting the temporal signal evolution in such studies, it is important to note that the hemodynamic response reflects brain activity in an indirect way and hence has a lag of about 5-6 s relative to neural activity (Liao et al., 2002). Four our data, this implies that mechanisms involved in influencing Gestalt perception appear to start operating already around 7-8 s before the actual stimulus presentation. Although seemingly long, this delay is consistent with previous studies investigating the prestimulus predictors of the motor and non-motor voluntary decisions, which found that the outcome of a decision was encoded in the brain activity 10 s before participants became aware of it (Soon et al., 2008, 2013). This long delay was interpreted to reflect preparatory mechanisms of higher control areas leading to the final decision. Similarly, we conclude that higher-level perceptual and general alertness mechanisms, that seem to shape the subsequent percept in favor of an illusory Gestalt, may already start around 7-8 s before stimulus onset, preparing an individual for their subsequent perceptual decision.

### Correlation vs. causality

Looking at the current results, one would be tempted to assume a causal link between prestimulus insula activity and subsequent perceptual outcome. In general, two conditions should be met before attributing a causal connection between brain activity and behavior (Bielczyk et al., 2018). That is, brain activity should precede said behavior and direct manipulation of brain activity or network inference methods should reveal an immediate behavioral effect. While the former is true for our data, the latter is yet to be confirmed using techniques more focused on causal inference, such as Granger Causality (Granger, 1969; Weisz et al., 2014) or Dynamic Causal Modeling (Friston et al., 2003). We emphasize that until more evidence is provided, causal interpretations in our and similar studies require caution.

### Conclusion

Taken together, our findings show an involvement of the bilateral dorsal insula in biasing the interpretation of a bistable Gestalt stimulus in favor of the holistic percept, and also suggest a possible contribution of the percept-selective IPS. They therefore imply an interplay between alertness-related and perception-related areas in determining the success of perceptual grouping. These areas could be a starting point for future studies of prestimulus network-level interactions between different brain areas leading to Gestalt formation.

## 5. CReDIT authorship contribution statement

**M. Wilding:** Conceptualization, Methodology, Software, Formal analysis, Investigation, Writing – original draft, Visualization,

**C. Körner:** Investigation, Resources, Writing – review & editing,

**A. Ischebeck:** Writing – review & editing, Supervision.

**N. Zaretskaya:** Conceptualization, Methodology, Software, Investigation, Resources, Writing – review & editing, Supervision, Funding acquisition.

## 6. Data availability

The data analyzed in this study (with the exception of raw volumetric anatomical and functional MRI data) are available from the corresponding author upon request. The current data protection policy of the University of Graz prohibits raw MRI data sharing.

## 7. Code availability

The code utilized in this study is available from the corresponding author upon request.

## 8. Acknowledgements

We thank Thomas Zussner, Adam Coates and Erik Fink for their data acquisition support, and Ana Arsenovic for fruitful discussions. Additionally, we thank the lab of molecular biotechnology from the TU Graz for their help. This work was funded by the BioTechMed-Graz Young Research Group Grant to N.Z.

## Notes

**Funding:** This work was supported by the BioTechMed-Graz Young Researcher Groups Grant Program.

### Competing Interest Statement

The authors have declared no competing interest.

