## Supplementary material for "Increased insula activity precedes the formation of subjective Gestalt"

Supplementary figure S1

Average activity for both conditions for summed poststimulus timepoints in all regions of interest with their corresponding time courses. For poststimulus comparison, the central 6 timepoints were summed (indicated by the grey area). Error bars represent the SEM (±) over subjects. Significant differences are indicated by an asterisk (*p<.05).

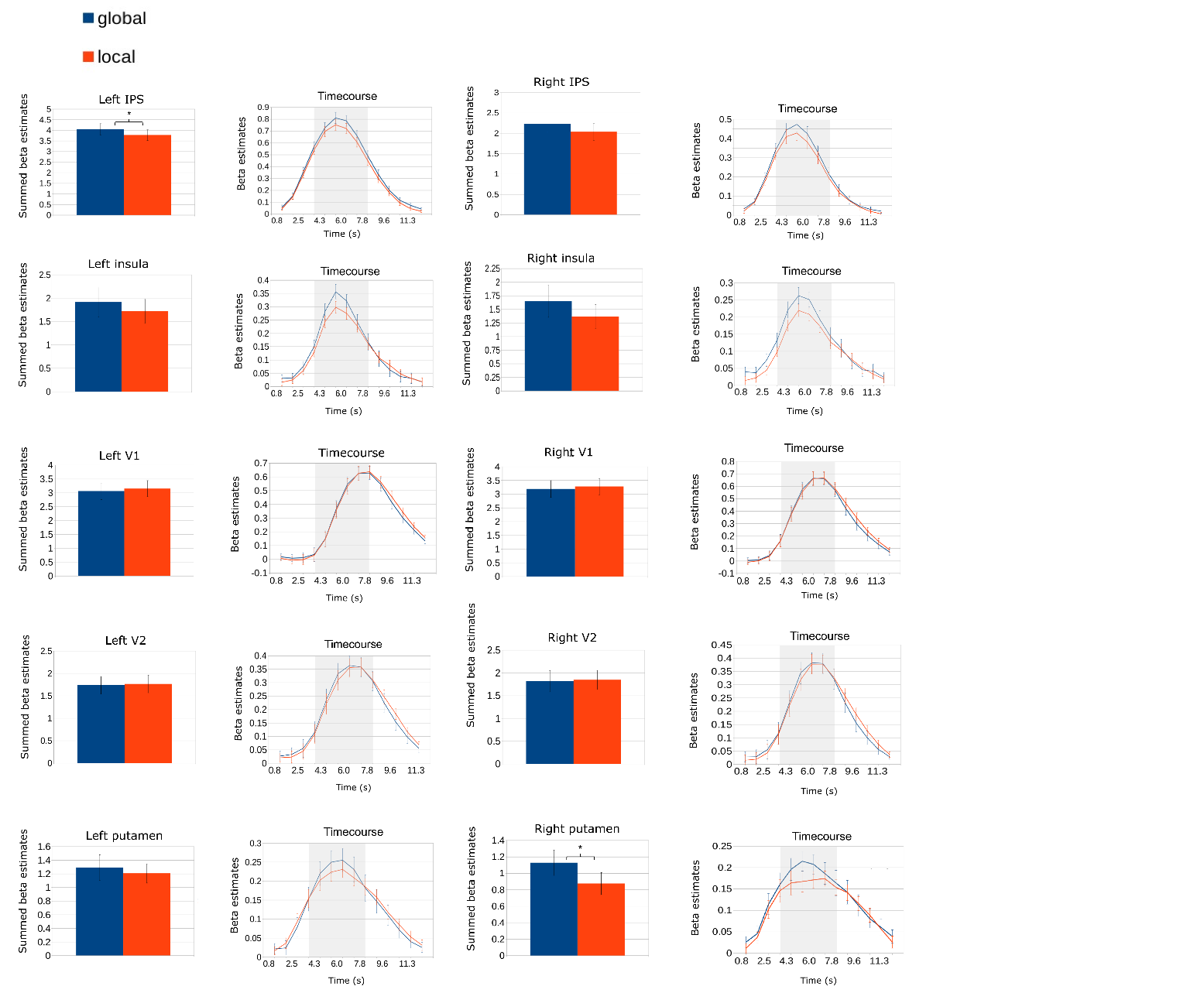

Supplementary figure S2

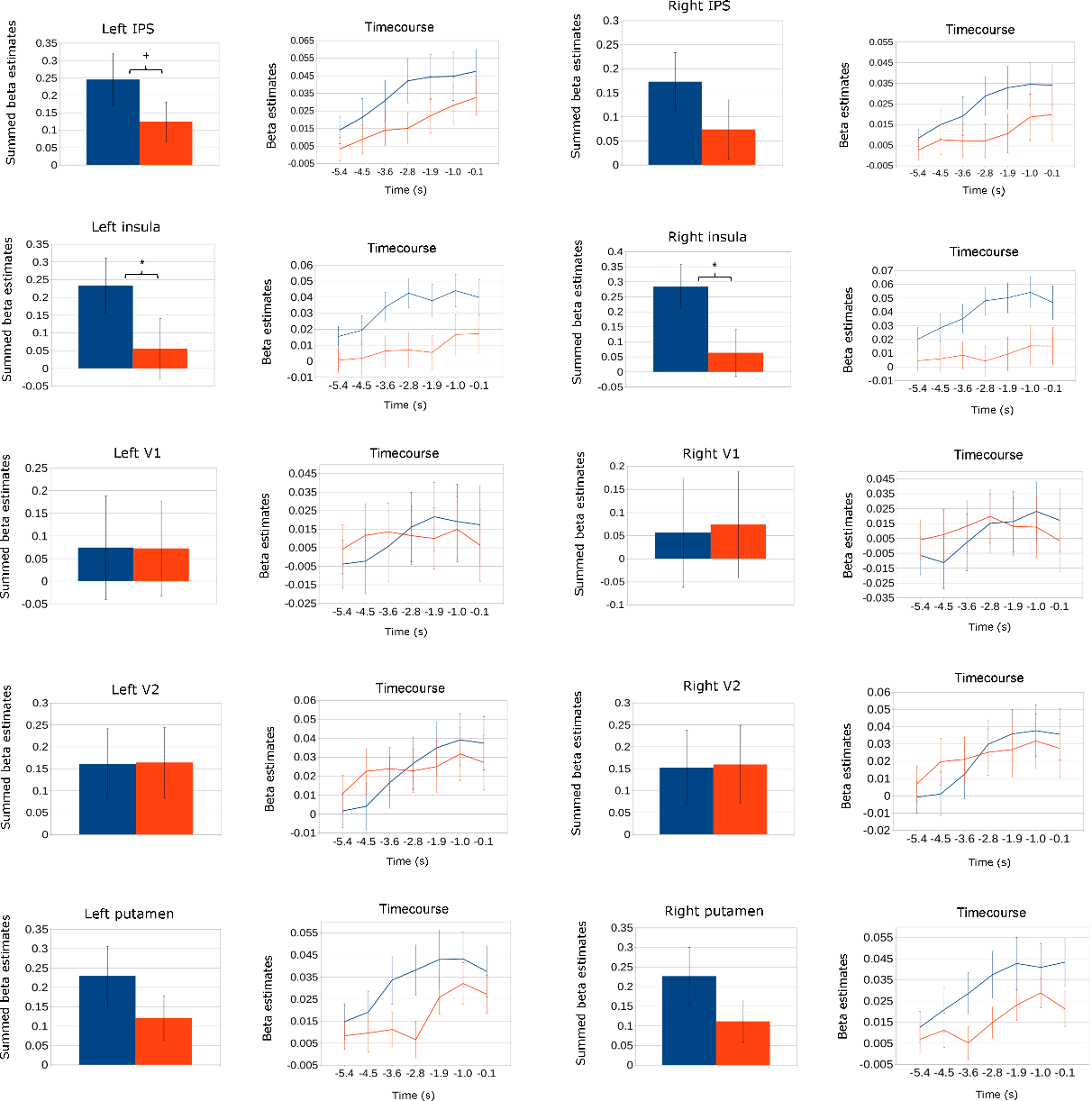
Average activity for both conditions for summed prestimulus timepoints in all regions of interest with their corresponding time courses. Prestimulus values were derived from the FIR-GLM model within a time window of -5.4 to -0.12 s, relative to the stimulus onset. Error bars represent the SEM (±) over subjects. Significant differences are indicated by an asterisk (*p<.05). A trend (p<.1) is indicated by a “+” sign.
